# 2’,3’-cAMP treatment mimics stress molecular response in *Arabidopsis thaliana*

**DOI:** 10.1101/2021.07.12.452129

**Authors:** Monika Chodasiewicz, Olga Kerber, Michal Gorka, Juan C. Moreno, Israel Maruri-Lopez, Romina I. Minen, Arun Sampathkumar, Andrew D. L. Nelson, Aleksandra Skirycz

## Abstract

The role of the RNA degradation product 2’,3’-cyclic adenosine monophosphate (2’,3’-cAMP) is poorly understood. Recent studies have identified 2’,3’-cAMP in plant material and determined its role in stress signaling. The level of 2’,3’-cAMP increases upon wounding, dark, and heat, and 2’,3’-cAMP by binding to an RNA-binding protein, Rbp47b, promotes stress granule (SG) assembly. To gain further mechanistic insight into 2’,3’-cAMP function, we used a multi-omics approach combining transcriptomics, metabolomics, and proteomics to dissect *Arabidopsis* response to 2’,3’-cAMP treatment. We demonstrated that 2’,3’-cAMP is metabolized into adenosine, suggesting that the well-known cyclic nucleotide–adenosine pathway from human cells might also exist in plants. Transcriptomic analysis revealed only minor overlap between 2’,3’-cAMP-and adenosine-treated plants, suggesting that these molecules act through independent mechanisms. Treatment with 2’,3’-cAMP changed the levels of hundreds of transcripts, proteins, and metabolites, many previously associated with plant stress responses including protein and RNA degradation products, glucosinolates, chaperones and SG components. Finally, we demonstrated that 2’,3’-cAMP treatment influences the movement of processing bodies, supporting the role of 2’,3’-cAMP in the formation and motility of membraneless organelles.

## Introduction

To cope with a fluctuating environment, living organisms developed signaling mechanisms to respond rapidly and acclimate to changing conditions. Signaling cascades comprise diverse protein and small molecule players, which act through a series of timely and spatially spaced interactions to regulate the activity, localization, and aggregation of downstream targets driving physiological alterations (Catozzi et al., 2016). Cyclic nucleotides are a group of important and evolutionarily conserved signaling small molecules (Manganiello and Degerman, 1999). In human cells, 3’,5’-cyclic adenosine monophosphate (cAMP) acts as a second messenger downstream of adrenaline and glucagon but upstream of sugar and lipid metabolism. In contrast with 3’,5’-cAMP, its positional isomer—2’,3’-cAMP—has received considerably less attention. In fact, 2’,3’-cAMP was only discovered in 2009 in a biological material (Ren et al., 2009). Further functional studies showed that it is a product of 3’→5’ RNA degradation (Thompson et al., 1994) and thus accumulates under conditions characterized by excessive mRNA decay, such as tissue injury (Jackson et al., 2009; Verrier et al., 2012; Van Damme et al., 2014). Even though this attribute is also true for other 2’,3’-cNMPs, 2’,3’-cAMP is the most abundant, mostly due to the presence of the mRNA poly(A) tail. In animal cells, high levels of cellular 2’,3’-cAMP are considered toxic and have been linked to mitochondrial dysfunction (Azarashvili et al., 2009). Furthermore, animal cells can efficiently metabolize 2’,3’-cAMP, first to 2’-AMP *via* the activity of the 2’,3’-cyclic nucleotide-3’-phosphodiesterase and subsequently to adenosine (Jackson et al., 2009; Jackson, 2016). Given that adenosine exhibits health-promoting properties (Jackson et al., 2009), the conversion of 2’,3’-cAMP to adenosine is proposed as a switch from a toxic to a non-toxic cellular environment.

In both animal and plant cells, cellular levels of 2’,3’-cAMP increase under stress treatments, such as wounding (Van Damme et al., 2014) or heat and dark conditions (Kosmacz et al., 2018). Moreover, 2’,3’-cAMP interacts with Rbp47b (Kosmacz et al., 2018; Kosmacz and Skirycz, 2020), an important protein for stress granule (SG) formation (Kosmacz et al., 2019). These findings support the role of 2’,3’-cAMP in stress signaling and regulation. SGs are non-membranous organelles formed in response to stress (Sorenson and Bailey-Serres, 2014; Gutierrez-Beltran et al., 2015; Jang et al., 2020). In addition to the core proteins required for the SG assembly and maintenance, such as protein and RNA chaperones, SG sequester a variety of metabolic enzymes and regulators. SGs are tightly linked to a different non-membrane organelle called processing bodies (PBs) (Kedersha et al., 2005), both of which are involved in the regulation of the fate of mRNA by affecting storage, degradation, and translation of mRNA (Chantarachot and Bailey-Serres, 2018).

To gain further mechanistic insight into 2’,3’-cAMP function, we used transcriptomics, metabolomics, and proteomics to dissect the *Arabidopsis* response to supplementation with a permeable analogue of 2’,3’-cAMP, Br-2’,3’-cAMP. We used the Br-analogue because without this modification, cyclic nucleotides cannot efficiently enter cells (Robison et al., 1965). The Br-derivatives of both 3’,5’-cAMP and 2’,3’-cAMP have been successfully used to study cAMP signaling and regulation in plants (Alqurashi et al., 2016; Kosmacz et al., 2018) and animals (Huising et al., 2007). Br-3’,5’-cAMP activates protein kinase A as efficiently as 3’,5’-cAMP, demonstrating that the addition of a bromide group does not interfere with cAMP physiological activity (Yao et al., 2015). Similarly, both exogenously supplied 2’,3’-cAMP and Br-2’,3’-cAMP induce exosome production in human carcinoma cells; however, the effective concentration of Br-2’,3’-cAMP is 10-fold lower than that of a 2’,3’-cAMP, which is attributed to the difference in uptake (Ludwig et al., 2020).We are the first to evaluate the response to Br-2’,3’-cAMP at the molecular level by combining complementary transcriptomics, proteomics and metabolomics experiments. Our data revealed the following: (i) Br-2’,3’-cAMP is taken up by plants, where it is metabolized to Br-adenosine; (ii) Br-2’,3’-cAMP triggers major responses at the transcript, protein, and metabolite levels, bearing a known stress signature; and (iii) treatment with Br-2’,3’-cAMP affects the abundance of key SG proteins and induces PB movement.

## Results

### Treatment with 2’,3’-cAMP leads to the accumulation of stress-responsive metabolites

To characterize plant response to the accumulation of 2’,3’-cAMP, we performed feeding experiments by treating *Arabidopsis* seedlings growing in liquid cultures with 1 µM of Br-2’,3’-cAMP, a membrane-permeable analogue of 2’,3’-cAMP. Samples were harvested after 15 and 30 min and 1, 6, and 24 h of treatment with either mock solution (control samples) or Br-2’,3’-cAMP (treated samples; **Figure 1A**). We confirmed the uptake of Br-2’,3’-cAMP by using liquid chromatography–mass spectrometry (LC–MS)-based metabolomics (**Figure 1B**). After 15 min, the samples already showed sufficient Br-2’,3’-cAMP to detect its accumulation in treated seedlings. The level peaked at 30 min, dropped sharply at 1 h, and decreased further at 6 h. Strikingly, no Br-2’,3’-cAMP was detected in the samples taken at 24 h, suggesting a rapid turn-over of the compound. In support of an active conversion of 2’,3’-cAMP into adenosine, a decrease in Br-2’,3’-cAMP was accompanied by an increase in the Br-adenosine levels. Again, only traces of Br-adenosine were detected in the samples taken at 24 h, suggesting further decay.

**Figure 1.**
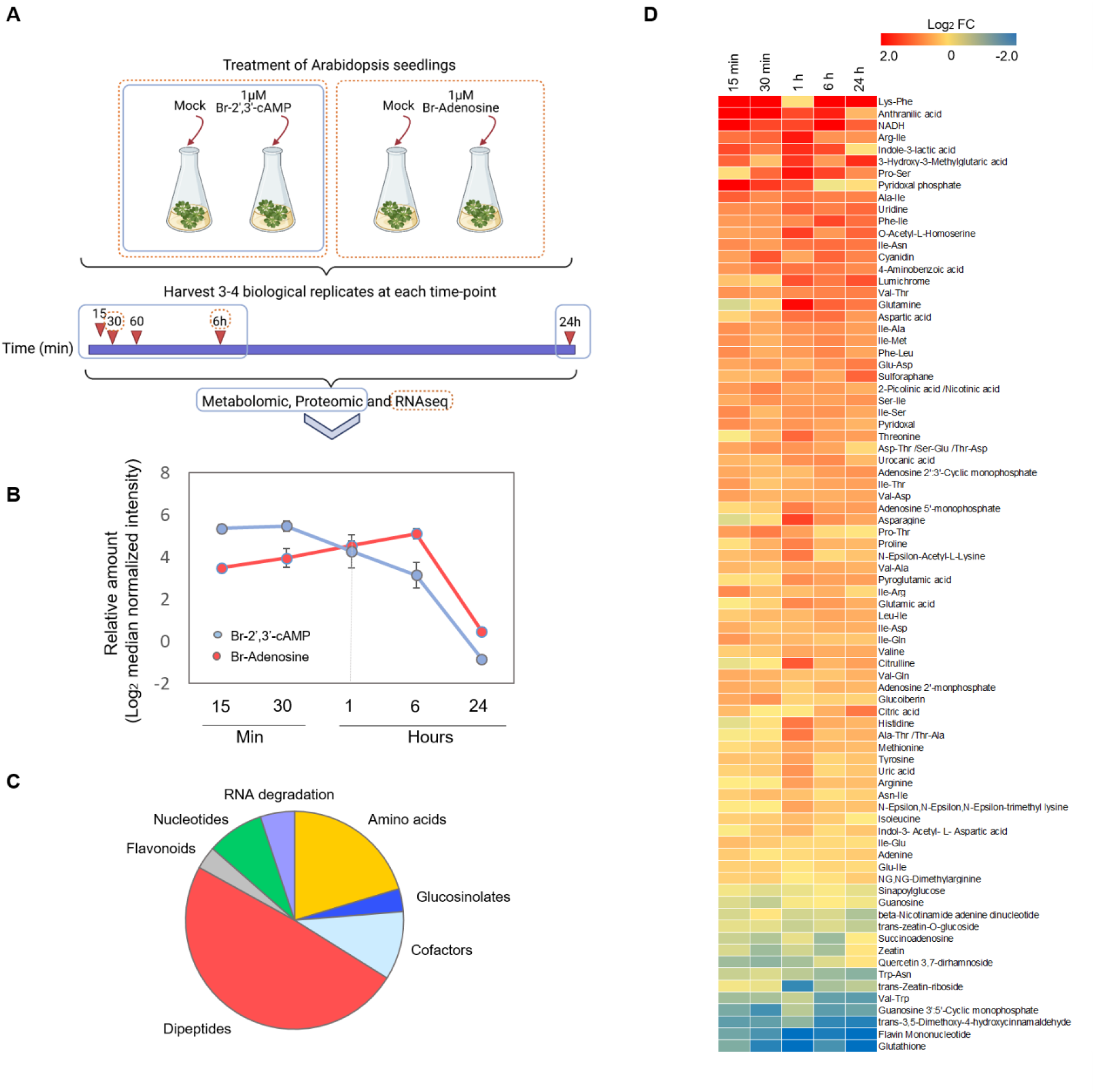
Br-2’,3’-cAMP induces stress-responsive changes at the metabolome level. **A**. Schematic representation of the experimental design. *Arabidopsis* wild-type plants were treated with mock, 1 μM Br-2’,3’-cAMP, or 1 µM Br-adenosine (for RNAseq analysis–orange rectangle/dashed lines). A total of 3–4 biological samples were collected at five time points for proteomic and metabolomic analyses (blue rectangle): 15 and 30 min, 1, 6, and 24 h, and only two time points at 15 min and 6 h for the transcriptome analysis (orange dashed line circles). Samples were extracted and prepared for proteomic, metabolomic, and RNAseq analyses. Data were analyzed with a focus on significant 2’,3’-cAMP-induced changes. **B**. Change in the amount of Br-2’,3’-cAMP and Br-adenosine levels in plants treated with 1 µM Br-2’,3’-cAMP. Data are presented as mean of log_2_ median normalized intensity. Error bars represent Standard Deviation n=4. **C**. The groups of metabolites significantly changed upon 2’,3’-cAMP treatment (two-way ANOVA, p-value FDR corrected ≤ 0.05). **D**. Heat map representing overall significant changes in metabolite levels after treatment with Br-2’,3’-cAMP. Data are presented as Log_2_ FC (two-way ANOVA, p-value FDR corrected ≤ 0.05).

In addition to Br-2’,3’-cAMP and Br-adenosine, metabolomic analysis (**Supplementary Table 1, Supplementary Table 1A**) identified 142 primary and specialized metabolites, 80 of which were significantly affected by the treatment with Br-2’,3’-cAMP (two-way analysis of variance [ANOVA], p-value FDR corrected ≤ 0.05; **Figure 1C, D**), and 68 of which were upregulated (**Figure 1D**). The accumulation of amino acids and proteogenic dipeptides suggests that Br-2’,3’-cAMP induces protein degradation (**Figure 1C**). Autophagy-dependent accumulation of dipeptides has been reported in plants subjected to heat and dark conditions (Thirumalaikumar et al., 2020). Autophagy is also a known source of amino acids (Hirota et al., 2018). Among the upregulated metabolites, endogenous 2’,3’-cAMP, adenosine-2’-monophosphate, and a 2’,3’-cAMP degradation product were detected, in addition to uric acid, which is a product of AMP catabolism (Hauck et al., 2014). This finding suggests that Br-2’,3’-cAMP treatment induces RNA decay, which is a characteristic of stressful conditions (Kosmacz et al., 2018). In summary, 2’,3’-cAMP supplementation leads to major metabolic alterations, reminiscent of stress conditions associated with high protein and mRNA turnover rates.

### Transcriptome analysis revealed that Br-2’,3’-cAMP mimics a stress response

Based on the major differences observed in the metabolome, we expected that Br-2’,3’-cAMP treatment would also significantly affect the transcriptome. Given that metabolomic analysis revealed an accumulation of adenosine in response to Br-2’,3’-cAMP supplementation, we also investigated to what extent the Br-2’,3’-cAMP response is related to the Br-adenosine build-up. Hence, we performed transcriptional profiling at two time points, 30 min and 6 h, for the Br-2’,3’-cAMP and Br-adenosine treatments. In our analysis, differentially expressed genes (DEGs) were genes that were significantly upregulated or downregulated (p-value adj. ≤ 0.05) compared with mock-treated samples. We compared the two datasets, which we referred to as 2’,3’-cAMP (Br-2’,3’-cAMP vs. mock-treated) and adenosine (Br-adenosine vs. mock-treated) treatments. We observed high reproducibility between samples (Supplemental Figure 1), with both compounds significantly affected gene expression (**Figure 2A; Supplementary Tables 2 and 3**). Adenosine treatment resulted in 2322 and 2798 DEGs at 30 min and 6 h, respectively. From the 2322 DEGs measured at 30 min, 547 DEGs were also differentially expressed at 6 h. By contrast, 2’,3’-cAMP treatment resulted in 953 and 2959 DEGs at 30 min and 6 h, respectively. From the 953 DEGs measured at 30 min, 439 DEGs were also differentially expressed at 6 h.

**Figure 2.**
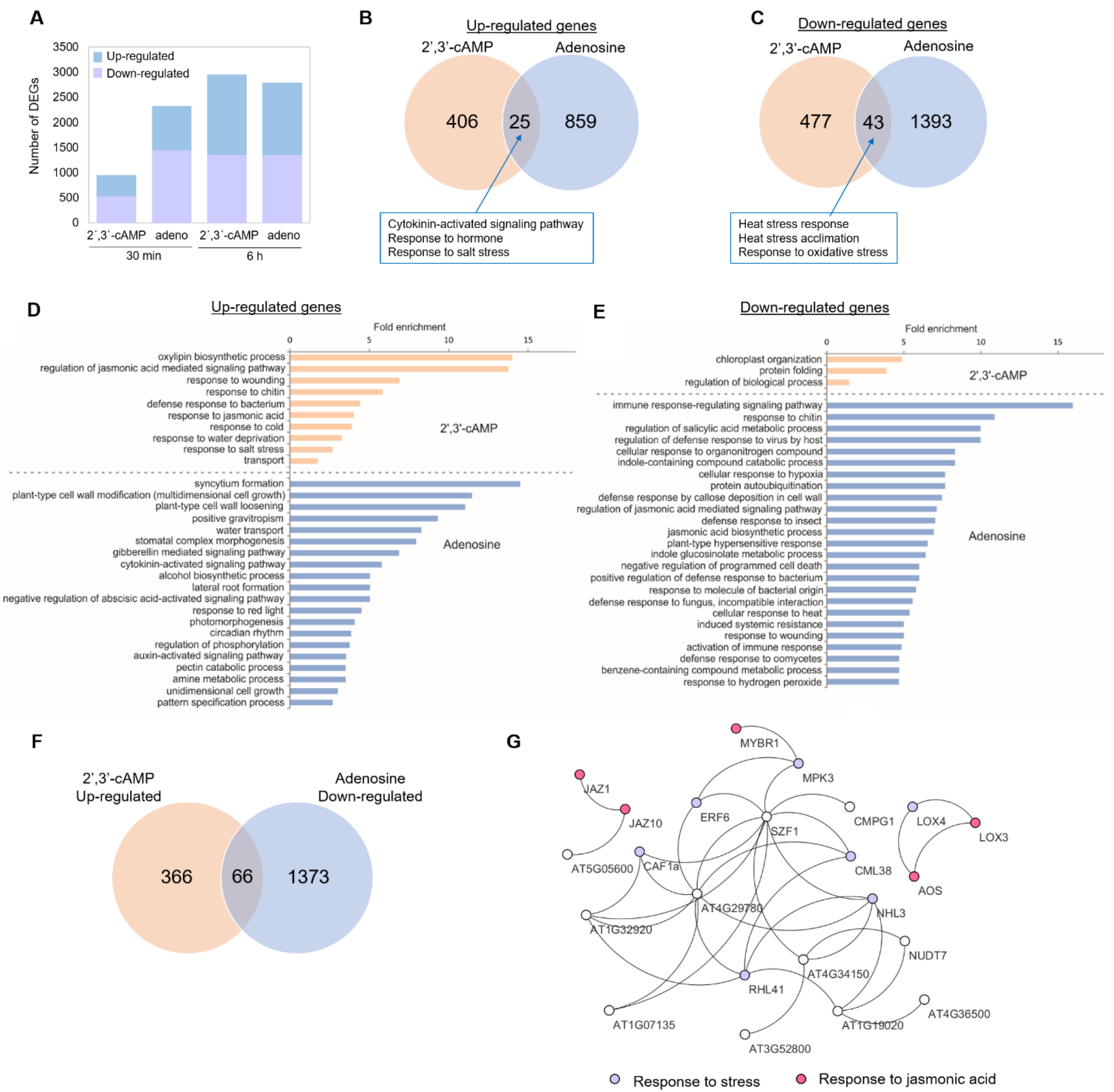
Differential gene expression analysis revealed major transcriptional reprogramming associated with 2’,3’-cAMP and adenosine. **A**. The number of genes found to be upregulated or downregulated after 30 min and 6 h of treatment. **B**. Venn diagram of all significantly upregulated genes after 30 min of 2’,3’-cAMP and adenosine treatment. **C**. Venn diagram of all significantly downregulated genes after 30 min of 2’,3’-cAMP and adenosine treatment. The numbers (**B, C**) correspond to DEGs identified as significantly changed compared with the control samples in each experiment. **D**. Overrepresentation of biological processes in the dataset of upregulated genes in 2’,3’-cAMP (light orange bars) and adenosine (light blue bars) experiments (30 min time-point). **E**. Overrepresentation of biological processes in the dataset of downregulated genes in 2’,3’-cAMP (light orange bars) and adenosine (light blue bars) experiments (30 min time-point). In (D–E), overrepresentation is shown as a significant fold enrichment based on the PANTHER Overrepresentation Test (Mi et al., 2017) by using Fisher’s exact test with FDR multiple correction (p-value ≤ 0.05) and *Arabidopsis thaliana* as the reference organism. **F**. Venn diagram showing an overlap between downregulated genes in adenosine treatment and upregulated genes in 2’,3’-cAMP treatment. **G**. A network of enriched genes was retrieved by the STRING database (Szklarczyk et al., 2017) but visualized in Cytoskape. Experimental and database evidence and a low confidence cut-off were used to visualize protein– protein interactions. Genes encoding for proteins involved in stress (violet) and response to JA (pink) are highlighted.

The comparison between DEGs identified in adenosine and 2’,3’-cAMP treatment at 30 min (**Figure 2B, C**) and 6 h (**Supplementary Figure 2**) revealed only minor overlap, demonstrating that most of the changes measured in response to 2’,3’-cAMP supplementation are specific to 2’,3’-cAMP (**Figure 2B, C**). An enrichment analysis of the different biological processes (Fisher’s exact test with an FDR correction, p-value 0.05) was performed using the PANTHER overrepresentation test (PANTHER13.1) with the GO ontology database (Mi et al., 2017). DEGs upregulated by 2’,3’-cAMP and adenosine at 30 min belonged to the groups “cytokinin-activated signaling pathway”, “hormonal response”, and “response to salt stress” (**Supplementary Table 4**). DEGs downregulated by both 2’,3’-cAMP and adenosine were enriched in genes involved in “heat stress response”, “acclimation”, and “response to oxidative stress” (**Supplementary Figure 2A, B and Supplementary Table 4**). 2’,3’-cAMP-specific DEGs induced at 30 min were enriched in transcripts associated with “oxylipin biosynthetic process”, “JA signaling pathways”, and “response to wounding” (**Figure 2D**; **Supplementary Figure 2A, B and Supplementary Table 5**), whereas 2’,3’-cAMP downregulated genes comprised multiple transcripts involved in “chloroplast organization” and “protein folding”, including multiple heat shock proteins and genes involved in the heat stress response (**Supplementary Figure 2A**). Approximately 30 min of adenosine treatment resulted in the upregulation of transcripts enriched in biological processes, such as “syncytium formation”, “cell wall modification”, and “hormonal signaling (gibberellin mediated signaling pathway)” (**Figure 2D**; **Supplementary Table 7**). Moreover, MapMan (Usadel et al., 2009) analysis (**Supplementary Figure 3**) revealed the over-representation of DNA-binding with one finger (Dof) transcription factors, which are involved in biotic stress response, synthesis of seed storage proteins, seed development, photosynthetic processes, and flowering (Wen et al., 2016). By contrast, “immune response”, “defense regulating pathways”, “indole-containing compound catabolic processes”, and “response to hypoxia” were highly enriched among the downregulated transcripts (**Figure 2E; Supplementary Table 6**). Interestingly, comparing 2’,3’-cAMP-upregulated genes and adenosine-downregulated genes revealed an overlap of 65 genes, mostly involved in stress response, specifically in “response to wounding and JA” (**Figure 2F, G**; **Supplementary Table 7**). This finding suggests that 2’,3’-cAMP and adenosine might have an antagonistic function in the cell in response to stress (e.g., wounding) but a synergistic function in the cytokinin-activated signaling pathway (upregulation) and response to heat (downregulation). In comparison to the samples induced at 30 min, the main GO categories enriched for the 2’,3’-cAMP-induced transcripts included “photosynthesis”, “hormonal response”, and “stress response”, whereas “response to oxidative stress”, and “metabolism” were included for adenosine (**Supplementary Figure 2A and Supplementary Table 8**). 2’,3’-cAMP-downregulated transcripts were enriched for “the RNA machinery and processing”, which again supported the role of 2’,3’-cAMP in the regulation of stress-related response (**Supplementary Figure 2B and Supplementary Table 9**). Finally, adenosine-downregulated transcripts were enriched for “hormonal response to gibberellin” and “hormonal transport”.

Because 2’,3’-cAMP was previously shown to play a role in the induction of SG formation (Kosmacz et al., 2018), we decided to check how many DEGs affected by 2’,3’-cAMP treatment contain the prion-like domain (PrLD), associated with the formation of liquid-liquid phase separation foci. We focused on the 30 min time-point, as in the past we could demonstrate that 30 min Br-2’, 3’-cAMP treatment is sufficient to induce SGs in the Arabidopsis seedlings (Kosmacz et al., 2018). Out of 953 DEGs (at 30 min), 29 encode proteins with a predicted PrLD (**Supplementary Figure 4**), including core component of SGs-Rbp47b. Moreover, proteins like ARF11 (Auxin responsive factor 11), PUM5 and PUM6 (pumilio genes), BHL3 (BEL1-like homeodomain 3) or TCP10 (TCP domain protein 10) are also on the list.

In summary, we demonstrated that 2’,3’-cAMP and adenosine induced a distinct set of genes, many previously associated with plant stress responses.

### Br-2’,3’-cAMP affects proteins associated with stress response and metabolism

To gain additional insight into the response to Br-2’,3’-cAMP treatment, we analyzed the changes in the abundance of proteins at all five time-points. Statistical analysis identified 472 differentially abundant proteins (two-way ANOVA, p-value FDR corrected ≤ 0.05; **Supplementary Table 1**). Subcellular localization analysis revealed an enrichment of plastid and cytosol localization among upregulated proteins, whereas nuclear and cytosol-localized among downregulated proteins. Functional analysis of upregulated proteins (PANTHER) identified functional categories, such as “amino acids, glucosinolates, nucleotide and pigment biosynthesis”, “photosynthesis”, and “auxin transport”. Downregulated proteins, analogous to downregulated transcripts, were enriched for proteins associated with protein folding, energy metabolism, response to heat, and translation (**Figure 3B**).

**Figure 3.**
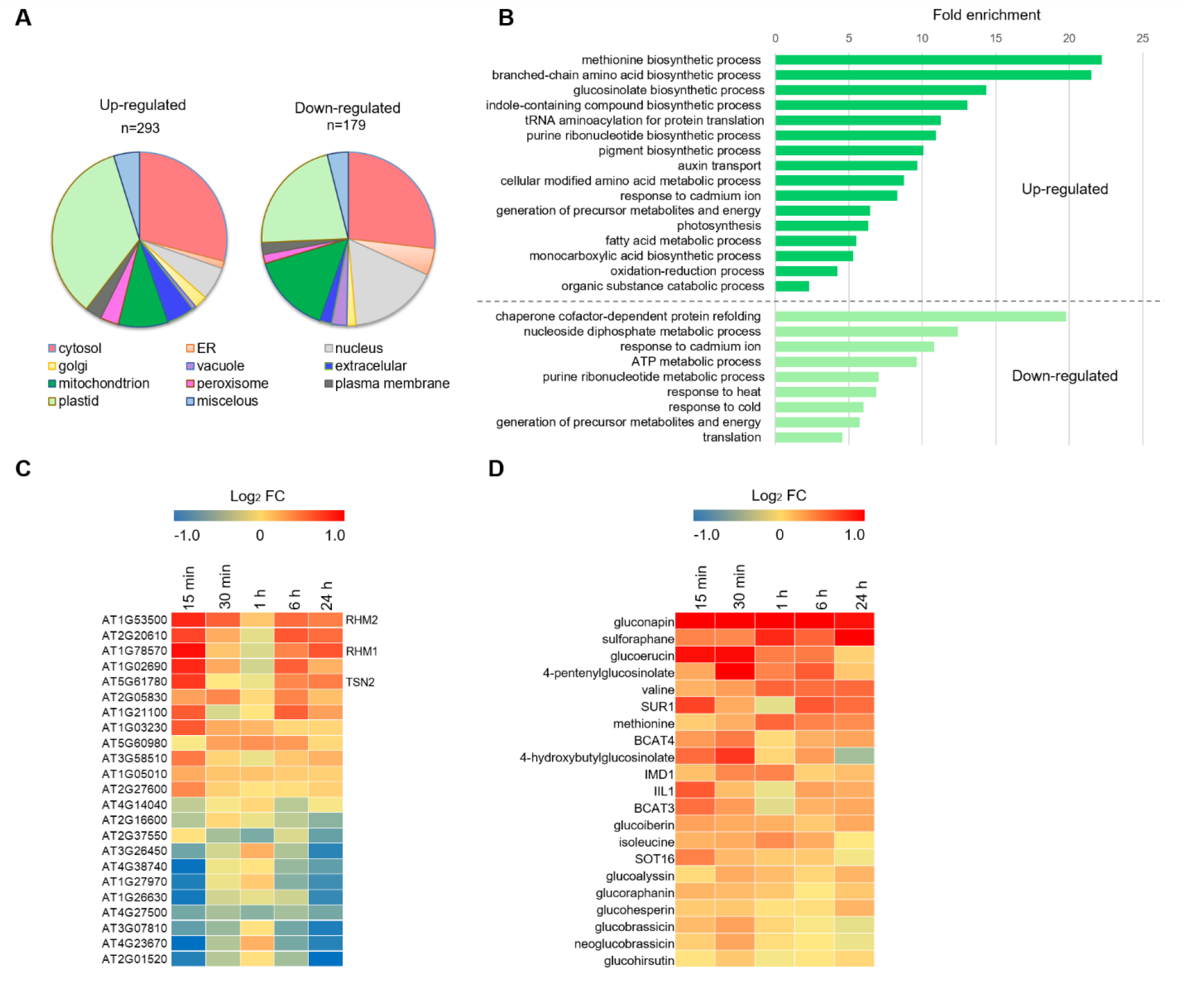
2’,3’-cAMP induces stress-related changes in the proteome of *Arabidopsis thaliana*. **A**. The cellular compartment distribution of the identified, significantly upregulated, and downregulated proteins after Br-2’,3’-cAMP treatment. Subcellular localizations for each protein were identified using the SUBA3 database (http://suba3.plantenergy.uwa.edu.au/). **B**. Enriched biological processes in the dataset of significantly upregulated (dark green) and downregulated (light green) proteins for Br-2’,3’-cAMP treatment. Overrepresentation is shown as a significant fold enrichment based on the PANTHER Overrepresentation Test (Mi et al., 2017) using Fisher’s exact test with FDR multiple correction (p-value ≤ 0.05) and *A. thaliana* as reference organism. **C**. Changes in the abundance of SG proteins are presented as heat maps. Significant changes were determined using a two-way ANOVA (p-value FDR corrected ≤ 0.05, *n* = 4 biological replicates). **D**. Heat map represents significantly accumulating putative aliphatic and indole glucosinolate compounds together with glucosinolate biosynthetic enzymes in response to Br-2’,3’-cAMP. Significant changes were determined using a two-way ANOVA (p-value FDR corrected ≤ 0.05, n = 4 biological replicates).

Because 2’,3’-cAMP accumulates in response to wounding (Van Damme et al., 2014) and Br-2’,3’-cAMP supplementation induces genes associated with jasmonate synthesis and signaling (see above), we were intrigued by the accumulation of glucosinolate biosynthetic enzymes. Glucosinolates are sulfur-containing specialized metabolites that contribute to plant defense against pests, and glucosinolate accumulation depends on jasmonate signaling (Mitreiter and Gigolashvili, 2021). To further investigate our observation, we searched for glucosinolates within the unknown features of our metabolomics dataset. Annotation based on a previous study (Wu et al., 2018) led to the identification of 13 putative aliphatic and indole glucosinolate compounds, 11 of which accumulated in response to the Br-2’,3’-cAMP treatment (two-way ANOVA, p-value ≤ 0.05; **Supplementary Table 1B, Figure 3D**). Whereas glucosinolate accumulation peaks at 15-30 min of Br-2’,3’-cAMP application, sulforaphane, a product of glucosinolate break-down, builds-up gradually and reaches its maximum after 24 h. Moreover, the list of 2’,3’-cAMP-responsive metabolites included glucosinolate precursor amino acids (methionine, valine, isoleucine), which ties with the accumulation of enzymes involved in branch-chain amino acid and methionine metabolism. However, it has to be noted that glucosinolate accumulation preceded the accumulation of amino acids. Finally, we investigated whether glucosinolate biosynthetic enzymes induced by Br-2’,3’-cAMP at the protein level are also induced at the transcript level, but this was not the case.

We also found that a crucial enzyme for aliphatic glucosinolate synthesis, (SUR1), which was previously found among proteins sequestered into cytosolic SG (Kosmacz et al., 2019), is also induced by Br-2’,3’-cAMP treatment. In addition to SUR1, 22 other proteins previously reported to localize to SG (**Figure 3C**; **Supplementary Table 1**) also respond to Br-2’,3’-cAMP supplementations, 12 upregulated and 11 downregulated. The list of Br-2’,3’-cAMP responsive proteins includes RHM2, RHM1 (Kosmacz et al., 2019), and TSN2 (Gutierrez-Beltran et al., 2015), providing additional evidence to link 2’,3’-cAMP and stress response at the SG level.

Finally, joined clustering analysis of protein and metabolite data delineated several patterns of accumulation (**Supplementary Figure 5, Supplementary Table 1C**). Notably, most proteins and metabolites, 257 of 563, were characterized by the largest change measured at the early 15 min and subsequently late 6 h and 24 h time-points, with more minor or no changes at 30 min and 1 h (**Supplementary Figure 5 and Supplementary Table 1C**). In comparison, Br-2’,3’-cAMP peaked at 30 min followed by a gradual decline, with no compound detected at 24 h. At first perplexing, we attribute this behavior to endogenous 2’,3’-cAMP. 2’,3’-cAMP levels were induced by the Br-2’,3’-cAMP treatment, with the highest accumulation measured at 6 h and 24 h. Among proteins that follow the pattern of Br-2’,3’-cAMP/2’,3’-cAMP accumulation are proteins located in the cluster 4 of 26 down-regulated proteins and metabolites (**Supplementary Figure 5A**) such as chaperones, proteasome subunit (PAG1) or GRF10 protein (14-3-3-like protein GF14 epsilon) from Brassionosteroid pathway.

### 2’,3’-cAMP induces movement of PBs

Given that SGs are highly connected with PBs and that 2’,3’-cAMP is closely related to RNA metabolism (being its degradation product), we investigated whether 2’,3’-cAMP can also affect PBs. The main function of PBs is translational repression and mRNA decay (Kedersha et al., 2005; Xu and Chua, 2011). However, both PBs and SG have continuous interactions (Anderson and Kedersha, 2009) and are often transiently linked to each other (Eisinger-Mathason et al., 2008; Buchan et al., 2012). In contrast with SGs, PBs are always present in the cell independently from stress conditions; however, stress may affect the dynamic of PBs. To evaluate this feature, we decided to focus on GFP-Decapping protein 1 (DCP1) protein, which is a well-known PB marker that was previously used for colocalization studies with SGs (Gutierrez-Beltran et al., 2015), and determined whether the treatment affects PB dynamics. Using confocal microscopy and aligning t-stags (**Figure 4A, B**), we followed the movement of PBs over time. Particle tracking showed that the application of Br-2’,3’-cAMP significantly induced displacement length (**Figure 4C**; **Supplementary Table 11**) and PB movement speed (**Figure 4D**) compared with those in the control samples (**Supplementary Table 12**). Our experiments indicate that 2’,3’-cAMP induces genes which contain PrLD but also affects proteins known to localize to SG/PBs. This could explain why the dynamic of PBs is affected which further suggests a role of 2’,3’-cAMP in the regulation of PB motility.

**Figure 4.**
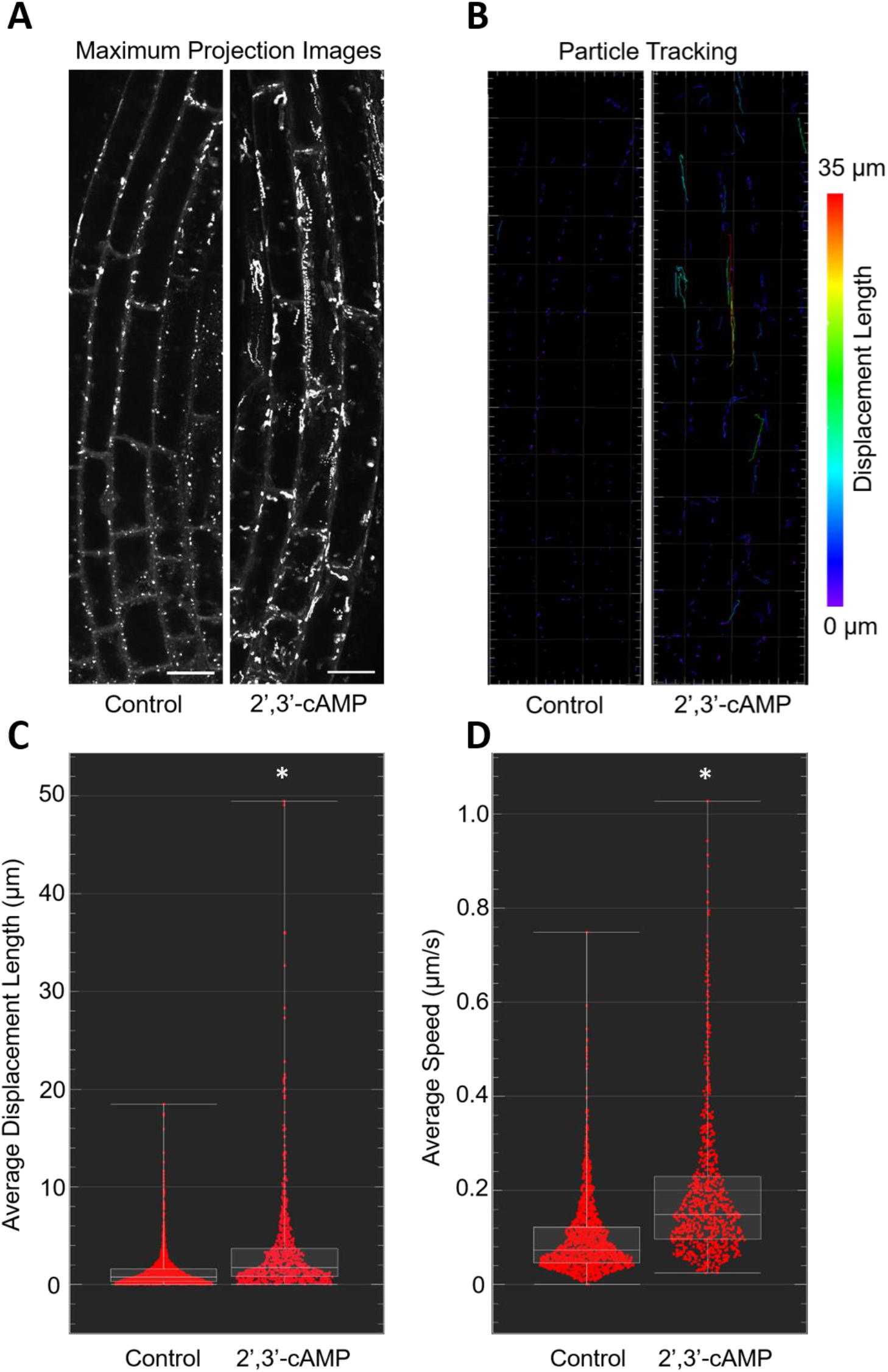
Br-2’,3’-cAMP induces motility of PBs. **A**. Maximum projection images from 61 time points collected from GFP-DCP1 *Arabidopsis* seedlings under control and Br-2’,3’-cAMP treatment. Scale bar = 10 µm. **B**. Particle tracking in the control and Br-2’,3’-cAMP -treated cells. Scale represents the color code for the distance of displacement for each PB. **C**. The average displacement length of the PB particles in the cell is expressed in µm. **D**. The average speed of PB movement in the control and Br-2’,3’-cAMP -treated seedlings. Speed is expressed in µm/s. For **C** and **D**, control *n* = 1583, 2’,3’-cAMP *n* = 898. Asterisk in C and D indicates significant differences defined by Student’s T-test, p-value ≤0.05.

## Discussion

Given that 2’,3’-cAMP has been detected in plants (Pabst et al., 2010), responds to stress conditions (Van Damme et al., 2014), and is a facilitator of SG formation (Kosmacz et al., 2018), characterizing its function further at different molecular levels is necessary. Our multi-omics approach (Moreno et al., 2021) is a valuable strategy to gain further insights at different molecular levels. To our knowledge, this study is the first time-resolved study exploring the comparison of cellular responses in plants at the metabolome, proteome, and transcriptome levels upon membrane-permeable 2’,3’-cAMP (Br-2’,3’-cAMP) treatment.

In animal cells, 2’,3’-cAMP is metabolized to adenosine. Our metabolomic analysis revealed that after 30 min of treatment in plants, Br-2’,3’-cAMP decreases and Br-adenosine increases (**Figure 1A**), suggesting the existence of the 2’,3’-cAMP adenosine salvage pathway in plants. Notably, these results are in line with previous findings where it was shown tha putative 2’,3’-cNMP cyclic phosphodiesterase, an enzyme involved in digestion of cNMPs, exists in *A. thaliana* (Genschik et al., 1997). This study further supports the finding that 2’,3’-cAMP is metabolized in plant cells and is a part of the adenosine salvage pathway that has only been described to exist in mammalian cells (Jackson et al., 2009). Interestingly, in plants, the nucleoside transporter (ENT) is important in regulating the levels of 2’,3’-cAMP and adenosine in the cell. Transgenic *Arabidopsis* plants with either low or high expression of the ENT are characterized by concomitant changes in the 2’,3’-cAMP and adenosine levels (Bernard et al., 2011). This finding again suggests that the concentration of those two compounds is tightly regulated by the presence of adenosine pathway components.

To distinguish between 2’,3’-cAMP and adenosine-specific changes, we first evaluated transcriptional response to 2’,3’-cAMP and adenosine (Br-adenosine). Transcriptomic data revealed that even though the genes that were upregulated by 2’,3’-cAMP and downregulated by adenosine overlapped, overall changes caused by those two compounds were significantly different. This result points toward distinct functions for 2’,3’-cAMP and adenosine. Genes and proteins affected by 2’,3’-cAMP are involved in several biological processes but are especially enriched in known stress markers, for example, ATG10 (AT3G07525), which is involved in the formation of autophagy vesicles; JAZ1 (AT1G19180) and JAZ10 (AT5G13220), which are central components of jasmonate signaling; critical stress kinase—MPK3 (AT3G45640), transcription factors, such as MYB44 (AT5G67300); and markers of the heat response such as HSP70 (AT3G12580) and MBFC1 (AT3G24500). Furthermore, metabolomics analyses has revealed the accumulation of known stress markers, such as glucosinolates, RNA-degradation products, and proteogenic dipeptides (**Figure 1C**) (Doppler et al., 2019; Thirumalaikumar et al., 2020; Moreno et al., 2021). The latter is especially interesting given the novel regulatory roles of dipeptides, such as in the regulation of enzymes of central carbon metabolism. Dipeptide feeding affects metabolic fluxes, leading to changes in metabolite pools impacting plant growth under oxidative stress (Moreno et al., 2021) and diauxic shift transition in yeast (Luzarowski et al., 2021). Hence, we speculate that the 2’,3’-cAMP-related accumulation of dipeptides contributes to the 2’,3’-cAMP response.

2’,3’-cAMP binds to the RNA-binding motifs (RRM) of Rbp47b, an SG assembly protein. In line with the binding data, 2’,3’-cAMP treatment facilitates SG formation (Kosmacz et al., 2018). RRM domains are not restricted to the Rbp47b protein. In fact, RRM is one of the most abundant protein domains in eukaryotes; it is also present in proteins associated with different facets of RNA metabolism, including RNA sequestration to non-membrane aggregates, such as SG. Among the 31 of the recently predicted 2’,3’-cAMP targets, seven contain the RRM domain (Zuhlke et al., 2021), including a plastidial protein CP29, which is a component of plastidial SGs (Chodasiewicz et al., 2020). Here, we demonstrated that 2’,3’-cAMP, in addition to binding to the core SG proteins, affects the abundance of proteins that sequester into SGs in response to heat condition (Kosmacz et al., 2019), such as tudor staphylococcal nuclease (TSN) (Gutierrez-Beltran et al., 2015). Notably, TSN is also found in PBs; in addition to its scaffolding role, TSN is also involved in mRNA decapping (Gutierrez-Beltran et al., 2015). Intriguingly, we showed that 2’,3’-cAMP increases the motility of PBs. PB movement depends on myosin (Steffens et al., 2014) and on interactions with SGs. Various components are known to be exchanged between those two foci (Kedersha et al., 2005); therefore, any effect on SGs may directly affect PBs.

In summary, 2’,3’-cAMP treatment affects the levels of hundreds of transcripts, proteins, and metabolites, many of which have been previously associated with plant stress response (**Figure 5**). The response is rapid and specific, occurring within the first 15–30 min of the 2’,3’-cAMP supplementation. Future works will focus on further characterization of the downstream responses, including changes in PB motility and dipeptide accumulation.

**Figure 5.**
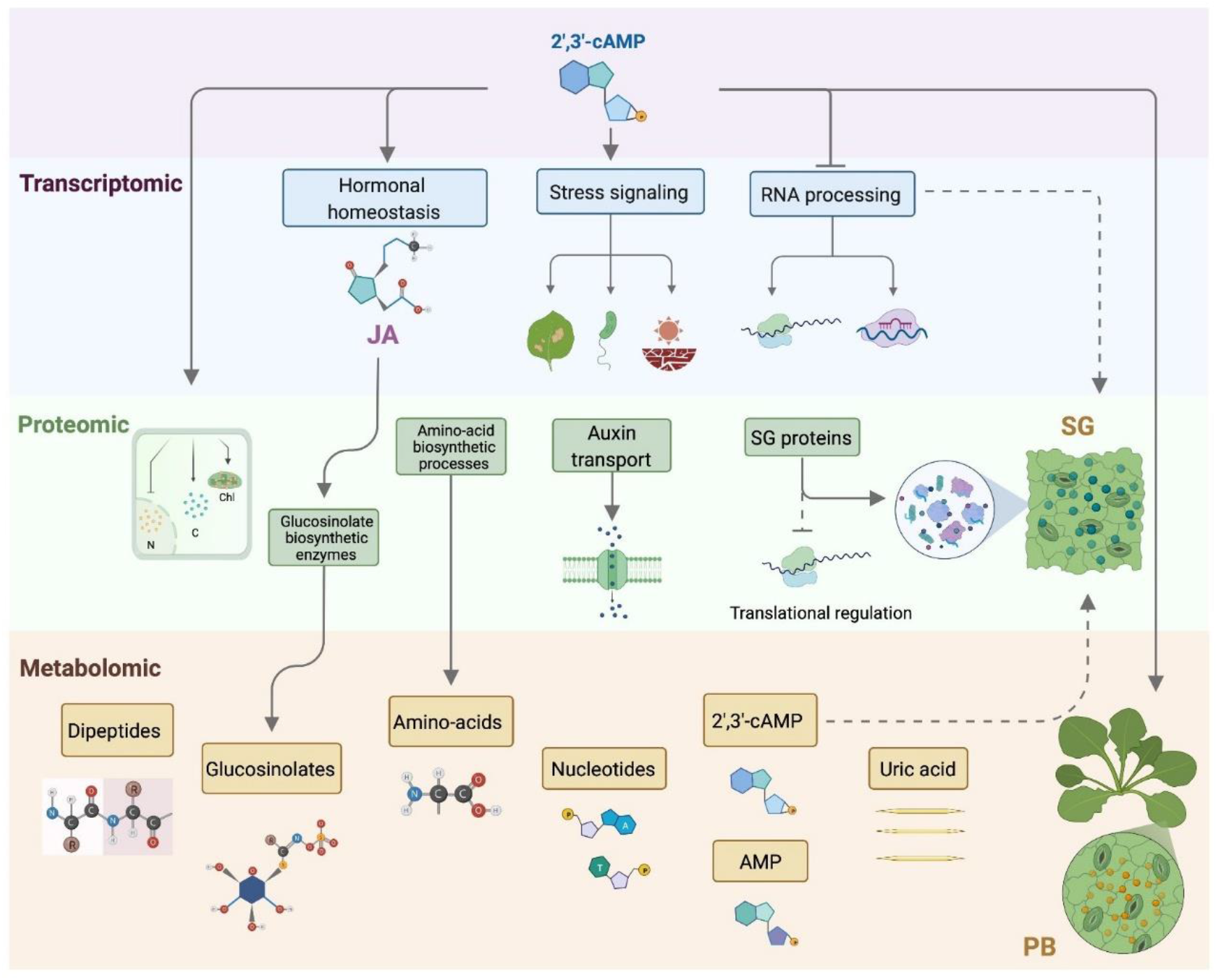
2’,3’-cAMP treatment resembles stress responses. Representative scheme of the effects of 2’,3’-cAMP (Purple panel) application in *A. thaliana* at transcriptomic, proteomic, and metabolomics levels. (Blue panel) Transcriptomic. Differential gene expression analysis showed that 2’,3’-cAMP upregulation of genes involved in Jasmonic acid (JA) homeostasis and stress responses (e.g. wounding, defense to bacteria, and water deprivation); whereas, those genes involved in RNA machinery and processing were downregulated during 2’,3’-cAMP treatment. Proteomic data (Green panel) revealed that 2’,3’-cAMP triggered stress-related changes in the *A. thaliana* proteome. Briefly, accumulation of proteins involved in amino acid biosynthetic pathways and auxin transport were increased upon 2’,3’-cAMP treatment, in contrast to translational machinery, which was downregulated. 2’,3’-cAMP also lead to accumulation of key SGs proteins and induced PBs movement. Metabolites (Orange panel) such as dipeptides, amino acids, nucleotides, and RNA-degradation products showed to accumulate under 2’,3’-cAMP. Arrows indicate up-regulation and bars depict down-regulation. Dotted lines denotes associated process.

## Materials and Methods

### Plant growth conditions and feeding experiment

*A. thaliana* Col-0 seedlings were grown in liquid MS medium (Murashige and Skoog, 1962) supplied with 1% sucrose in continuous light. The medium was changed after 7 days, and treatment with 1 µM Br-2’,3’-cAMP (Biolog Life Science Institute) was performed after 3 days (at 10^th^ day). Approximately 1 µM was used based on low to mid µM concentrations of endogenous 2’,3’-cAMP measured previously in Arabidopsis native lysate (Kosmacz et al., 2018). As the control, the seedlings were treated with water (in which the compound was dissolved). The seedlings were then harvested after 15 and 30 min and 1, 6, and 24 h of treatment, rapidly dried on paper, and frozen in liquid nitrogen. Dry plant material was obtained by dry vacuum for 3 days. Experiment using the same conditions with 1 µM Br-adenosine (SIGMA-Aldrich) was performed for transcriptome analysis.

### Metabolite and protein extraction

The protocol for the extraction of molecules was adjusted from Salem et al. (2017). Using 10 mg of dried tissue powder, macromolecules were extracted using a methyl tert-butyl ether (MTBE)/methanol/water solvent system, which separates the molecules into pellet (proteins), organic (lipids), and aqueous phases (primary and secondary metabolites). Equal volumes of the collected fractions were dried using a centrifugal evaporator and stored at 80 °C before metabolomic and proteomic analyses.

### LC–MS metabolomics for secondary metabolite identification

The dried aqueous phase was measured using ultra-performance liquid chromatography coupled to an exactive mass spectrometer (Thermo Fisher Scientific, Bremen, Germany) in positive and negative ionization modes. The method was reproduced from Giavalisco et al. (2011). However, the spectra were recorded in full-scan mode only. Data processing was performed using REFINER MS 10.5 (GeneData; http://www.genedata.com) and included peak detection, chemical noise subtraction, retention-time (RT) alignment, and integration of isotopic peaks into peak clusters. Metabolite features were annotated using an in-house reference compound library allowing 10 ppm m/z and 0.1 min RT deviations. The library is composed of authentic chemical standards, which were run using the method described above. Obtained chromatograms were used to extract information on the main adduct and RT. Moreover, the library comprises phenylpropanoids, flavonols, and glucosinolates, which were annotated based on the fragmentation and elemental formula information as described by (Wu et al., 2018), using identical instruments and methods.

### LC–MS/MS for proteins and data analysis

Protein pellets formed in the MTBE-based extraction method were solubilized in 100 μl of urea– thiourea buffer (6 M urea, 2 M thiourea in 40 mM ammonium bicarbonate). Protein content was determined using a Bradford assay (Carl Roth, Karlsruhe, Germany). Approximately 40 µg of protein was treated with 5 mM of dithiothreitol for 30 min at RT, followed by cysteine alkylation with 15 mM iodoacetamide for 20 min at RT in the dark. Enzymatic digestion of proteins using LysC/Trypsin Mix (Promega, Fitchburg, WI) was then performed according to the technical manual. After digestion, samples were acidified with trifluoroacetic acid (TFA) to pH < 2. Peptides were desalted using C18 Empore® extraction discs (3M, Maplewood, MN) STAGE tips (Rappsilber et al., 2003) and dried to approximately 4 µl using a centrifugal evaporator. Samples were stored at 80 °C until measurement. Dried peptides were solubilized in loading buffer (2% ACN, 0.2% TFA), and an equivalent of 0.8–1.0 µg of peptides was separated using a reversed-phase column and analyzed on a Q-Exactive Plus or Q-Exactive HF spectrometer (Thermo Fisher Scientific). MaxQuant version 1.6.0.16 (Cox and Mann, 2008) and its built-in search engine Andromeda (Cox et al., 2011) were used to analyze the raw proteomic data. For protein annotation, the *A. thaliana* TAIR10 annotations (Arabidopsis TAIR database Version 10, The Arabidopsis Information Resource, www.Arabidopsis.org [updated in December 2017]) combined with the search engine Andromeda were used. The search also included a contaminant database that can be found in any MaxQuant installation folder or directly downloaded from the developer’s official website. Contaminants and decoy hits were removed from each dataset. The settings for MaxQuant analysis were set as follows: trypsin and lysine were selected as digesting enzymes, two missed cleavages were allowed, fixed modification was set to carbamidomethylation (cysteine), and oxidation of methionine was set as a variable modification. Spectra were also searched against a decoy database of the *A. thaliana* proteome, and results were filtered to obtain an FDR below 1% on the protein level. The “label-free quantification” and “match between runs” options were selected. A minimum peptide length of six amino acids was used. The quantification was performed for proteins with a minimum of one unique and one razor peptide. Known contaminants, such as keratins, were removed from further analysis. Furthermore, at least two unique peptides were required per protein group. Label-free quantification (LFQ) intensities were used in all analyses performed in this manuscript.

### RNA extraction and RNAseq analysis

An RNA extraction kit (Macherey-Negel) was used to extract total RNA from 10 mg of lyophilized tissue, followed by quality assessment by a Bioanalyzer RNA 6000 nano (Agilent). RNAseq analysis was performed by Lexogen GmbH (Lexogen, Vienna, Austria) using QuantSeq 3′-mRNA library preparation and QuantSeq 3′-UTR NextSeq SR75 sequencing. An integrated data analysis was performed by Lexogen (STAR aligner), where the reads were mapped against the *A. thaliana* reference genome (Araport11), read-counts were determined, and differential expression was computed using DESeq2 in R (Love et al., 2014). To perform RNAseq analysis, two biological replicates for two time points of 30 min and 6 h were used in each experiment. MapMan software (Usadel et al., 2009) was used to visualize perturbations in gene expression.

### Statistical analysis

GeneData-derived raw metabolite intensities were normalized to the median intensity of all mass features detected in a given chromatogram. MaxQuant-derived LFQ intensities were used for further analysis. Both metabolite and protein data were subjected to log_2_ transformation prior to two-way ANOVA analysis implemented in the MeV software (Howe et al., 2011) using treatment (treated vs. untreated) and time (15 min, 30 min, 1 h, 6 h, 24 h) as variables. The obtained p-values were subjected to FDR correction to select metabolites and proteins significantly affected by the treatment. The Software MeV (Howe et al., 2011) was used to obtain heat maps. Differential gene expression was computed using DESeq2 in R (Love et al., 2014). The obtained p-values were FDR corrected. For the proteomic and transcriptomic analyses, a fold enrichment analysis was performed using the PANTHER overrepresentation test (PANTHER13.1) based on Fisher’s exact test with FDR correction p-value ≤ 0.05. Subcellular localization analyses were retrieved from the SUBA3 database (Tanz et al., 2013). Clustering of proteomic and metabolomics data was performed also using MeV Software.

### Prion like domain analysis

PrLD analysis was performed using PLAAC software, where amino acid sequences of all significant DEGs from 30 min of Br-2’,3’-cAMP treatment were uploaded. Data are present in the **Supplementary Table 10**.

### Data deposition

The mass spectrometry proteomics data have been deposited to the ProteomeXchange Consortium via the PRIDE (Perez-Riverol et al., 2016) partner repository with the dataset identifier PXD028365. RNAseq raw data have been deposited to NCBI repository with the number PRJNA769461.

### Processing body dynamic assessment under confocal microscope

*Arabidopsis* seeds expressing GFP-tagged DCP1 (Gutierrez-Beltran et al., 2015), the PB marker, were kindly provided by Dr. Emilio Gutierrez-Beltran. Plants were grown for 5–7 days on MS media supplied with 1% sucrose. On the day of the experiment, seedlings were moved to an Eppendorf tube and incubated either with water (control) or with 100 µM Br-2’,3’-cAMP for 30 min. After incubation, seedlings were observed under a confocal microscope (Leica TCS SP8) using the x, y, t function. GFP was excited using a 488 nm laser, and emission was obtained between 500 and 600 nm. Images were collected every 2 s for a total of 69 images, which were used for video assembly. The calculations of PB displacement and displacement speed were done using the IMARIS software (https://imaris.oxinst.com/).

## Author contribution

MC, OK, and JCM performed the feeding experiments for Br-2’,3’-cAMP and Br-adenosine experiments; MC and OK prepared samples for metabolomics, transcriptomics, and proteomics; MG run and annotated the proteomic samples and deposited the data to PRIDE. Aleksandra S performed the annotation of metabolomic data and analysis; MC, ADLN, and Aleksandra S analyzed transcriptomic and proteomic data; IM-L performed data mining and prepared summary figure; MC performed confocal microscopy for PBs; RIM provided help and organized supplementary material; AS performed measurements on PB dynamics; Aleksandra S and MC conceived the project; ADLN submitted RNAseq raw data to NCBI depository, MC wrote the paper with the input from Aleksandra S, JCM and ADLN. All authors contributed to the final version of this manuscript.

## Supplementary Figures

**Supplemental Figure 1**. Clustering of transcriptomic profiles between replicates of treated and untreated seedlings. **A**. PCA plot of between sample variations. **B**. Sample distance matrix demonstrating reproducibility between replicates.

**Supplemental Figure 2**. Overrepresentation of the biological process in a set of up- and downregulated specific genes in 2’,3’-cAMP (orange bars) and adenosine experiments (blue bars). Overrepresentation is shown as a significant fold enrichment based on the PANTHER Overrepresentation Test (Mi et al., 2017).

**Supplemental Figure 3**. MapMan representation of transcriptional perturbations in 30 min time point for Br-2’,3’-cAMP (upper panel) and 30 min of Br-adenosine treatment. Schematic overview of different processes generated in MapMan depicting hormonal, signaling, and stress response perturbations. Significantly affected DEGs were used for each treatment. The red square represents the most affected processes/components of biological processes.

**Supplemental Figure 4**. Transcriptom-wide analysis identified 29 DEGs with PrLDs. Graph represents DEGs with PrLD, organized by the length. Aa – amino acids. Supplementary Table 10.

**Supplemental Figure 5**. Co-expression clustering between proteins and metabolites that are down-or up-regulated by Br-2’,3’-cAMP treatment. Data re provided in Supplementary Table 1C.

## References

Alqurashi M, Gehring C, Marondedze C (2016) Changes in the Arabidopsis thaliana Proteome Implicate cAMP in Biotic and Abiotic Stress Responses and Changes in Energy Metabolism. Int J Mol Sci 17

Anderson P, Kedersha N (2009) RNA granules: post-transcriptional and epigenetic modulators of gene expression. Nat Rev Mol Cell Biol 10: 430–436

Azarashvili T, Krestinina O, Galvita A, Grachev D, Baburina Y, Stricker R, Evtodienko Y, Reiser G (2009) Ca2+-dependent permeability transition regulation in rat brain mitochondria by 2’,3’-cyclic nucleotides and 2’,3’-cyclic nucleotide 3’-phosphodiesterase. Am J Physiol Cell Physiol 296: C1428–1439

Bernard C, Traub M, Kunz HH, Hach S, Trentmann O, Mohlmann T (2011) Equilibrative nucleoside transporter 1 (ENT1) is critical for pollen germination and vegetative growth in Arabidopsis. J Exp Bot 62: 4627–4637

Buchan JR, Capaldi AP, Parker R (2012) TOR-tured yeast find a new way to stand the heat. Mol Cell 47: 155–157

Catozzi S, Di-Bella JP, Ventura AC, Sepulchre JA (2016) Signaling cascades transmit information downstream and upstream but unlikely simultaneously. BMC Syst Biol 10: 84

Chantarachot T, Bailey-Serres J (2018) Polysomes, Stress Granules, and Processing Bodies: A Dynamic Triumvirate Controlling Cytoplasmic mRNA Fate and Function. Plant Physiol 176: 254–269

Chodasiewicz M, Sokolowska EM, Nelson-Dittrich AC, Masiuk A, Beltran JCM, Nelson ADL, Skirycz A (2020) Identification and Characterization of the Heat-Induced Plastidial Stress Granules Reveal New Insight Into Arabidopsis Stress Response. Front Plant Sci 11: 595792

Cox J, Mann M (2008) MaxQuant enables high peptide identification rates, individualized p.p.b.-range mass accuracies and proteome-wide protein quantification. Nature Biotechnology 26: 1367–1372

Cox J, Neuhauser N, Michalski A, Scheltema RA, Olsen JV, Mann M (2011) Andromeda: A Peptide Search Engine Integrated into the MaxQuant Environment. Journal of Proteome Research 10: 1794–1805

Doppler M, Kluger B, Bueschl C, Steiner B, Buerstmayr H, Lemmens M, Krska R, Adam G, Schuhmacher R (2019) Stable Isotope-Assisted Plant Metabolomics: Investigation of Phenylalanine-Related Metabolic Response in Wheat Upon Treatment With the Fusarium Virulence Factor Deoxynivalenol. Front Plant Sci 10: 1137

Eisinger-Mathason TS, Andrade J, Groehler AL, Clark DE, Muratore-Schroeder TL, Pasic L, Smith JA, Shabanowitz J, Hunt DF, Macara IG, Lannigan DA (2008) Codependent functions of RSK2 and the apoptosis-promoting factor TIA-1 in stress granule assembly and cell survival. Mol Cell 31: 722–736

Genschik P, Hall J, Filipowicz W (1997) Cloning and characterization of the Arabidopsis cyclic phosphodiesterase which hydrolyzes ADP-ribose 1’’,2’’-cyclic phosphate and nucleoside 2’,3’-cyclic phosphates. J Biol Chem 272: 13211–13219

Gutierrez-Beltran E, Moschou PN, Smertenko AP, Bozhkov PV (2015) Tudor Staphylococcal Nuclease Links Formation of Stress Granules and Processing Bodies with mRNA Catabolism in Arabidopsis. Plant Cell 27: 926–943

Hauck OK, Scharnberg J, Escobar NM, Wanner G, Giavalisco P, Witte CP (2014) Uric acid accumulation in an Arabidopsis urate oxidase mutant impairs seedling establishment by blocking peroxisome maintenance. Plant Cell 26: 3090–3100

Hirota T, Izumi M, Wada S, Makino A, Ishida H (2018) Vacuolar Protein Degradation via Autophagy Provides Substrates to Amino Acid Catabolic Pathways as an Adaptive Response to Sugar Starvation in Arabidopsis thaliana. Plant Cell Physiol 59: 1363–1376

Howe EA, Sinha R, Schlauch D, Quackenbush J (2011) RNA-Seq analysis in MeV. Bioinformatics 27: 3209–3210

Huising MO, van der Aa LM, Metz JR, de Fatima Mazon A, Kemenade BM, Flik G (2007) Corticotropin-releasing factor (CRF) and CRF-binding protein expression in and release from the head kidney of common carp: evolutionary conservation of the adrenal CRF system. J Endocrinol 193: 349–357

Jackson EK (2016) Discovery and Roles of 2’,3’-cAMP in Biological Systems. Handb Exp Pharmacol

Jackson EK, Ren J, Mi Z (2009) Extracellular 2’,3’-cAMP is a source of adenosine. J Biol Chem 284: 33097–33106

Jang GJ, Jang JC, Wu SH (2020) Dynamics and Functions of Stress Granules and Processing Bodies in Plants. Plants (Basel) 9

Kedersha N, Stoecklin G, Ayodele M, Yacono P, Lykke-Andersen J, Fritzler MJ, Scheuner D, Kaufman RJ, Golan DE, Anderson P (2005) Stress granules and processing bodies are dynamically linked sites of mRNP remodeling. J Cell Biol 169: 871–884

Kosmacz M, Gorka M, Schmidt S, Luzarowski M, Moreno JC, Szlachetko J, Leniak E, Sokolowska EM, Sofroni K, Schnittger A, Skirycz A (2019) Protein and metabolite composition of Arabidopsis stress granules. New Phytol

Kosmacz M, Luzarowski M, Kerber O, Leniak E, Gutierrez-Beltran E, Moreno JC, Gorka M, Szlachetko J, Veyel D, Graf A, Skirycz A (2018) Interaction of 2’,3’-cAMP with Rbp47b Plays a Role in Stress Granule Formation. Plant Physiol 177: 411–421

Kosmacz M, Luzarowski M, Kerber O, Leniak E, Gutierrez-Beltran E, Moreno JC, Gorka M, Szlachetko J, Veyel D, Graf A, Skirycz A (2018) Interaction of 2 ‘,3 ‘-cAMP with Rbp47b Plays a Role in Stress Granule Formation. Plant Physiol 177: 411–421

Kosmacz M, Skirycz A (2020) The Isolation of Stress Granules From Plant Material. Curr Protoc Plant Biol 5: e20118

Love MI, Huber W, Anders S (2014) Moderated estimation of fold change and dispersion for RNA-seq data with DESeq2. Genome Biol 15: 550

Ludwig N, Yerneni SS, Menshikova EV, Gillespie DG, Jackson EK, Whiteside TL (2020) Simultaneous Inhibition of Glycolysis and Oxidative Phosphorylation Triggers a Multi-Fold Increase in Secretion of Exosomes: Possible Role of 2’3’-cAMP. Sci Rep 10: 6948

Luzarowski M, Vicente R, Kiselev A, Wagner M, Schlossarek D, Erban A, de Souza LP, Childs D, Wojciechowska I, Luzarowska U, Gorka M, Sokolowska EM, Kosmacz M, Moreno JC, Brzezinska A, Vegesna B, Kopka J, Fernie AR, Willmitzer L, Ewald JC, Skirycz A (2021) Global mapping of protein-metabolite interactions in Saccharomyces cerevisiae reveals that Ser-Leu dipeptide regulates phosphoglycerate kinase activity. Commun Biol 4: 181

Manganiello VC, Degerman E (1999) Cyclic nucleotide phosphodiesterases (PDEs): diverse regulators of cyclic nucleotide signals and inviting molecular targets for novel therapeutic agents. Thromb Haemost 82: 407–411

Mi H, Huang X, Muruganujan A, Tang H, Mills C, Kang D, Thomas PD (2017) PANTHER version 11: expanded annotation data from Gene Ontology and Reactome pathways, and data analysis tool enhancements. Nucleic Acids Res 45: D183–D189

Mitreiter S, Gigolashvili T (2021) Regulation of glucosinolate biosynthesis. J Exp Bot 72: 70–91

Moreno JC, Martinez-Jaime S, Kosmacz M, Sokolowska EM, Schulz P, Fischer A, Luzarowska U, Havaux M, Skirycz A (2021) A Multi-OMICs Approach Sheds Light on the Higher Yield Phenotype and Enhanced Abiotic Stress Tolerance in Tobacco Lines Expressing the Carrot lycopene β-cyclase1 Gene. Frontiers in Plant Science 12

Moreno JC, Rojas BE, Vicente R, Gorka M, Matz T, Chodasiewicz M, Peralta-Ariza JS, Zhang Y, Alseekh S, Childs D, Luzarowski M, Nikoloski Z, Zarivach R, Walther D, Hartman MD, Figueroa CM, Iglesias AA, Fernie AR, Skirycz A (2021) Tyr-Asp inhibition of glyceraldehyde 3-phosphate dehydrogenase affects plant redox metabolism. EMBO J: e106800

Murashige T, Skoog F (1962) A Revised Medium for Rapid Growth and Bio Assays with Tobacco Tissue Cultures. Physiologia Plantarum 15: 473–497

Pabst M, Grass J, Fischl R, Leonard R, Jin C, Hinterkorner G, Borth N, Altmann F (2010) Nucleotide and nucleotide sugar analysis by liquid chromatography-electrospray ionization-mass spectrometry on surface-conditioned porous graphitic carbon. Anal Chem 82: 9782–9788

Perez-Riverol Y, Xu QW, Wang R, Uszkoreit J, Griss J, Sanchez A, Reisinger F, Csordas A, Ternent T, Del-Toro N, Dianes JA, Eisenacher M, Hermjakob H, Vizcaino JA (2016) PRIDE Inspector Toolsuite: Moving Toward a Universal Visualization Tool for Proteomics Data Standard Formats and Quality Assessment of ProteomeXchange Datasets. Mol Cell Proteomics 15: 305–317

Rappsilber J, Ishihama Y, Mann M (2003) Stop and go extraction tips for matrix-assisted laser desorption/ionization, nanoelectrospray, and LC/MS sample pretreatment in proteomics. Anal Chem 75: 663–670

Ren J, Mi Z, Stewart NA, Jackson EK (2009) Identification and quantification of 2’,3’-cAMP release by the kidney. J Pharmacol Exp Ther 328: 855–865

Robison GA, Butcher RW, Oye I, Morgan HE, Sutherland EW (1965) The effect of epinephrine on adenosine 3’, 5’-phosphate levels in the isolated perfused rat heart. Mol Pharmacol 1: 168–177

Sorenson R, Bailey-Serres J (2014) Selective mRNA sequestration by OLIGOURIDYLATE-BINDING PROTEIN 1 contributes to translational control during hypoxia in Arabidopsis. Proc Natl Acad Sci U S A 111: 2373–2378

Steffens A, Jaegle B, Tresch A, Hulskamp M, Jakoby M (2014) Processing-body movement in Arabidopsis depends on an interaction between myosins and DECAPPING PROTEIN1. Plant Physiol 164: 1879–1892

Szklarczyk D, Morris JH, Cook H, Kuhn M, Wyder S, Simonovic M, Santos A, Doncheva NT, Roth A, Bork P, Jensen LJ, von Mering C (2017) The STRING database in 2017: quality-controlled protein-protein association networks, made broadly accessible. Nucleic Acids Res 45: D362–D368

Tanz SK, Castleden I, Hooper CM, Vacher M, Small I, Millar HA (2013) SUBA3: a database for integrating experimentation and prediction to define the SUBcellular location of proteins in Arabidopsis. Nucleic Acids Res 41: D1185–1191

Thirumalaikumar VP, Wagner M, Balazadeh S, Skirycz A (2020) Autophagy is responsible for the accumulation of proteogenic dipeptides in response to heat stress in Arabidopsis thaliana. FEBS J

Thompson JE, Venegas FD, Raines RT (1994) Energetics of catalysis by ribonucleases: fate of the 2’,3’-cyclic phosphodiester intermediate. Biochemistry 33: 7408–7414

Usadel B, Poree F, Nagel A, Lohse M, Czedik-Eysenberg A, Stitt M (2009) A guide to using MapMan to visualize and compare Omics data in plants: a case study in the crop species, Maize. Plant Cell Environ 32: 1211–1229

Van Damme T, Blancquaert D, Couturon P, Van Der Straeten D, Sandra P, Lynen F (2014) Wounding stress causes rapid increase in concentration of the naturally occurring 2’,3’-isomers of cyclic guanosine-and cyclic adenosine monophosphate (cGMP and cAMP) in plant tissues. Phytochemistry 103: 59–66

Verrier JD, Jackson TC, Bansal R, Kochanek PM, Puccio AM, Okonkwo DO, Jackson EK (2012) The brain in vivo expresses the 2’,3’-cAMP-adenosine pathway. J Neurochem 122: 115–125

Wen CL, Cheng Q, Zhao L, Mao A, Yang J, Yu S, Weng Y, Xu Y (2016) Identification and characterisation of Dof transcription factors in the cucumber genome. Sci Rep 6: 23072

Wu S, Tohge T, Cuadros-Inostroza A, Tong H, Tenenboim H, Kooke R, Meret M, Keurentjes JB, Nikoloski Z, Fernie AR, Willmitzer L, Brotman Y (2018) Mapping the Arabidopsis Metabolic Landscape by Untargeted Metabolomics at Different Environmental Conditions. Mol Plant 11: 118–134

Xu J, Chua NH (2011) Processing bodies and plant development. Curr Opin Plant Biol 14: 88–93

Yao Y, Cui X, Al-Ramahi I, Sun X, Li B, Hou J, Difiglia M, Palacino J, Wu ZY, Ma L, Botas J, Lu B (2015) A striatal-enriched intronic GPCR modulates huntingtin levels and toxicity. Elife 4

